# Stress attenuation by the adrenergic-specific lncRNA NESPR prevents cell death in neuroblastoma cells

**DOI:** 10.1101/2025.04.07.647560

**Authors:** Louis Delhaye, Dries Rombaut, Eric de Bony, Eva D’haene, Tiago França Brazao, Lisa Depestel, Bieke Decaesteker, Stéphane Van Haver, Ramiro Martinez, Laudonia Lidia Dipalo, Julliete Roels, Kimberly Verniers, Nurten Yigit, Jasper Anckaert, Bjorn Menten, Sarah Vergult, Pieter Van Vlierberghe, Stephen S. Roberts, Giorgio Milazzo, Giovanni Perini, Takaomi Sanda, Frank Speleman, Sven Eyckerman, Pieter Mestdagh

## Abstract

Neuroblastoma is a pediatric cancer of the sympathetic nervous system characterized by heterogeneous cell states that mirror normal differentiation trajectories. Each state is governed by a core regulatory transcriptional circuitry that reinforces cell identity through an autoregulatory feedforward loop. We identified the long non-coding RNA NESPR as specifically expressed in adrenergic neuroblastoma cells. NESPR expression correlates with high-risk neuroblastoma clinical parameters and poor patient survival. *NESPR* is located within an insulated gene neighborhood alongside PHOX2B, a master transcription factor of the adrenergic identity. NESPR depletion reduced cell proliferation and increased caspase activity in neuroblastoma cell lines, and NESPR knockout in a neuroblastoma zebrafish model led to reduced tumor penetrance. Subcellular localization revealed NESPR to be a cytosolic long non-coding RNA, suggesting a *trans*-regulatory function. RNA-sequencing following NESPR depletion revealed a shift from an adrenergic to a mesenchymal cell state, due to proteotoxic stress-induced molecular reprogramming. These findings suggest NESPR as a regulator of neuroblastoma cell identity with potential therapeutic opportunities in high-risk neuroblastoma cases.

## INTRODUCTION

Neuroblastoma (NB) is a pediatric solid malignancy that originates during development of truncal neural crest cells of the sympathoadrenal (SA) lineage(1,2). NB patients are stratified in low, intermediate, and high-risk groups based on prognostic molecular markers, reflecting a wide spectrum of clinical outcomes. Low and intermediate NB risk groups have high survival rates, with tumors responding to treatment or exhibiting spontaneous regression. Despite extensive advances in understanding NB tumor biology, high-risk NB survival rates are merely 45-55% with only minor improvements over the past decades(3,4). Although high-risk NB is generally a mutationally silent tumor, segmental copy number variations (CNV) such as amplification of the *MYCN* proto-oncogene, 17q gain, 1p loss, and 11q loss occur more frequently, and are associated with poor patient survival(5). In addition to CNVs, telomerase or alternative lengthening of telomeres (ALT) activation, due to *MYCN*, *TERT*, or *ATRX* genetic abnormalities, and alterations of RAS and p53 pathways frequently accompany these oncogenic driver events(6).

NB tumors demonstrate intratumoral heterogeneity, consisting of cellular identities that distinguish different molecular and morphological phenotypes resembling different differentiation stages of the SA lineage(7–9). These identities are established by the collective transcriptional activity of a so-called core regulatory transcriptional circuitry (CRC). Each CRC consists of a specific set of master transcription factors (mTF) that drive the expression of its own mTFs members as well as their distinct target genes. This effectively generates an autoregulatory feedforward loop that delineates the cell identity based on CRC target gene expression. Two major cell identities are discerned: the (nor)adrenergic (ADRN) cell identity resembles cells committed to the SA lineage that express mTFs such as PHOX2B, GATA3, ISL1, HAND2, TBX2, SOX11, MEIS2, and ASCL1(8–13). In contrast, the mesenchymal (MES) identity resembles a less differentiated cell type with migratory neural crest-like properties(14). ADRN tumors express proliferative genes and exhibit an aggressive disease progression but also present a more therapy-prone phenotype. MES subtype cells generally manifest a slower progression yet with a migratory and more therapy-resistant phenotype(15,16). Depending on extra- or intratumoral triggers such as chemotherapy or genetic mutations, both subtypes can interconvert(17,18), making it imperative to understand both subtypes to advance towards effective multimodal treatment regimes.

The non-coding transcriptome accounts for the majority of all transcribed genes, most of which are long non-coding RNAs (lncRNAs)(19). Many lncRNAs show a pronounced tissue- or cell type-specific expression profile(20), making lncRNAs attractive for therapeutic or diagnostic applications. Indeed, a handful of lncRNAs have been implicated as either tumor suppressive or oncogenic in NB. For example, several *MYCN*-co-amplified lncRNAs have been shown to regulate MYCN levels in high-risk NB, while others are overexpressed in *MYCN*-amplified tumors(21–24). In contrast, lncRNAs expressed from the 6p22.3 susceptibility locus have tumor suppressive properties and are frequently silenced(25,26). Other lncRNAs, such as HAND2-AS1 (also known as DEIN), TBX2-AS1, and SILC1 are products of the high transcriptional activity that is inherently concentrated at CRC mTF genes and their super-enhancer (SE) regions(11,12,27). Modi et al. (28) recently assessed tissue specificity, regulation, and putative function of all expressed lncRNAs across a pan-pediatric tumor panel, including the TARGET and GMKF NB cohorts. NB patients demonstrated the most abundant and highest tissue-specific expression across all tumor entities. Integration of ADRN mTF DNA binding sites, chromatin interaction patterns, and differential expression established 300 ADRN-associated lncRNAs. Silencing of the top-ranking TBX2-AS1 resulted in impaired NB cell growth due to E2F1 target gene downregulation. These analyses highlight the currently untapped therapeutic potential that lncRNAs can have in combatting childhood cancers.

Here, we highlight LINC00682, which we renamed to NESPR (Neuroblastoma-specific non-coding RNA). We outline NESPR’s unique expression profile in ADRN NB tumors and cell lines, and characterize its locus in context of its protein-coding neighbor PHOX2B, a ADRN mTF. Moreover, molecular, cell viability, and survival phenotypes following NESPR depletion indicate NESPR attenuates cellular stress to prevent cell death in ADRN NB cells.

## MATERIAL AND METHODS

### Public databases and data resources

RNA Atlas(29) transcriptome data was retrieved from the R2 Genomics Analysis and Visualization platform (http://r2platform.com/rna_atlas). CCLE(30) transcriptome data (2019 release) was retrieved through the DepMap portal (https://depmap.org/portal). Expression data of 10011 primary tumors from 34 different cancer types was obtained through the TCGA data portal (https://portal.gdc.cancer.gov). Raw data and clinical metadata of the SEQC(31) NB patient cohort were retrieved from the Gene Expression Omnibus (GEO id: GSE62564), and expression data was processed as described below and mapped to the hg19 human reference genome assembly (GRCh37). Expression data and clinical metadata of the Westermann (R2 id ps_avgpres_nbsewester579_gencode19) NB patient cohort was retrieved from the R2 platform. DESeq2-normalized counts of the SAP differentiation trajectory were obtained from Van Haver et al. (32). Fetal neuroblast and adrenal cortex expression data were retrieved from De Preter et al. (33). Raw gene expression data for GIMEN (GEO id: GSM2664383) was obtained from Boeva et al. (9), and mapped to the hg19 human reference genome (GRCh37). H3K27ac ChIP-seq from IMR32, SK-N-BE(2)c, SHEP, and GIMEN were retrieved from Boeva et al. (9) (GEO id: GSE90683). SMC1 HiChIP in CLBGA (GEO id: GSM4042589) and Kelly (GEO id: GSM2756147) cell lines was obtained from Gartlgruber et al. (34) and Debruyne et al. (35), respectively. ADRN mTF ChIP-seq in SK-N-BE(2)c data was obtained from Durbin et al. (8) (GEO id: GSE94822). CAGE profiles for NB-1 (FF ontology id: 10539-107G8): and CHP-134 (FF ontology id: 10508-107D4) cell lines were obtained from the FANTOM5 consortium (https://fantom.gsc.riken.jp/5/). H3K4me1 and H3K4me3 ChIP-seq data from SK-N-BE(2)c cells were retrieved from Upton et al. (36) (GEO id: GSE138315). Nuclear (GEO id: GSM2400226) and cytoplasmic (GEO id: GSM2400225) fractionation RNA-seq data of SK-N-SH cells was obtained from the ENCODE consortium(37) (https://www.encodeproject.org). Th-MYCN RNA-seq data (GEO id: GSM3519713 to GSM3519716) was obtained from Dorstyn et al. (38). Tg(dβh:EGFP-MYCN) RNA-seq data was obtained from Nunes et al. (39). Other publicly available data resources are listed through the main text.

### Cell culture

#### Cell lines

SK-N-BE(2)c, NGP, IMR32, SH-SY5Y, and SHEP cell lines were cultured in RPMI 1640 supplemented with 10% FBS, 2 mM glutamine, and 30 U/mL Penicillin-Streptomycin. Cell cultures were maintained under 60-70% confluency and passaged twice a week. Cells lines were confirmed mycoplasma-free on a regular basis by a mycoplasma colorimetric detection assay on medium of 48 h-confluent cultures. STR genotyping was performed to validate cell line authenticity. Cells were cultured in designated incubators at 37°C with a humidified atmosphere containing 5% CO_2_.

#### Antisense oligonucleotide transfection

Intronic and exonic ASOs were designed with the LNCASO software (https://iomics.ugent.be/lncaso) with default settings and the hg19 human reference genome assembly (GRCh37). The splice-junction targeting ASO was designed manually. All ASOs were LNA-modified, and contained a GapmeR configuration with a phosphorothiorate backbone.

Cells were transfected with 100 nM ASO using lipofectamine 2000 according to the manufacturer’s instructions. ASO sequences are listed in Table S1.

### Cell survival and proliferation

#### IncuCyte live cell imaging

To conduct cell confluence measurements, 10 000 IMR32, 5000 SK-N-BE(2)c, 5000 NGP, or 8000 SHEP cells were seeded in a 96-well plate. After 24 h, cell adherence was assessed visually per cell line. Fully adhered cell lines were transfected with ASOs in randomized block over the middle of the plate as described before. Cell confluence was measured in an IncuCyte S3 Live-Cell instrument, capturing images per well every hour, after seeding up to 96 hours after ASO transfection. Experiments were repeated 6 times per cell line per ASO. IncuCyte measurements at 48 hours post-transfection were statistically compared between ASO-targeting ASO and NCA within each cell line using a non-parametric Wilcoxon rank-sum test was performed in R. Two-sided *P*-values were adjusted for multiple testing using the Benjamini-Hochberg method. Adjusted *P-*values ≤ 5% were considered statistically significant. Data visualization was performed with the ggplot2 (v3.4.4) R package.

#### Caspase-3/7 assays

To asses apoptosis induction, 10 000 IMR32, 5000 SK-N-BE(2)c, 5000 NGP, or 8000 SHEP cells were seeded in a 96-well plate. After 24 h, cell adherence was assessed visually per cell line. Fully adhered cell lines were transfected with ASOs in randomized block over the middle of the plate as described before. Forty-eight hours after transfection, caspase-3/7 activity was measured with a Caspase-Glo 3/7 Assay System according to the manufacturer’s instructions. Luminescent signal was detected with a plate reader using default settings. Contrasts were calculated by dividing the luminescent signal per replicate with the average luminescent signals of the NCA ASO transfections. Normalized luminescent signals were averaged over all replicates. Caspase-3/7 activity of each NESPR-targeting ASO compared to a NCA within each cell line was statistically assessed using the non-parametric Wilcoxon rank-sum test in R. Two-sided P-values were adjusted for multiple testing using the Benjamini-Hochberg method. Adjusted P-values ≤ 5% were considered statistically significant. Data visualization was performed with the ggplot2 (v3.4.4) R package.

### RNAscope

10 000 cells were seeded in 400 uL medium per chamber of a 8-chamber imaging slide and grown to 50-75% confluency. Next day, half the medium was replaced with 4% prewarmed PFA in PBS for a final concentration of 2% PFA. After 15 min incubation at room temperature, the total volume was replaced with 2% prewarmed PFA in PBS, and incubated for an additional 10 min. Slides were washes three times with PBS, and stored in PBS at 4°C until staining. Nuclei were stained using 1:1000-diluted DAPI for 15 min at room temperature. RNAscope staining was performed with custom NESPR-specific probes containing a Cy3 fluorophore, using the RNAscope Multiplex Fluorescent V2 assay according to the manufacturer’s instructions for adherent cells. *B. subtilis* DapB- and *H. sapiens* PPIB-targeting probes were included as negative and positive technical controls, respectively, for all experiments. Stained slides were stored at 4°C in PBS protected from light exposure until confocal imaging. All imaging was performed on a LSM880 Airyscan with a Plan-Apochromat 63x/1.4 oil DIC M27. For all images the Airyscan detector was operated in the super resolution mode of the FastAiryScen. Pixel reassignment and 2D wiener deconvolution was performed post-acquisition in ZEN Black 2.3 SP1. Processing of images was performed in ImageJ2 (v2.16.0/1.54p) implemented in Fiji. Processing of images was first performed on SHEP cells, and settings were then transferred to SK-N-BE(2)c and IMR32 cells. Thresholds to define nuclei based on DAPI signal was performed for each cell line separately by manual adjustment.

### RNA extraction, cDNA synthesis, and RT-qPCR

#### RNA extraction

Attached cells were washes once with PBS, trypsinized, and collected by centrifugation. After two more PBS washes, pellets were redissolved in 700 uL QIAzol, incubated for 15 min at room temperature, snap-frozen, and stored at −80°C until further processing. Samples were unthawed at room temperature and 140 uL 100% chloroform was added. After vigorous mixing and centrifugation at 12 000 x g for 15 min at 4°C, the upper aqueous phase was collected to which 600 uL 100% EtOH was added. RNA from the mixture was isolated with the miRNeasy Micro kit following the manufacturer’s instructions including a Dnase I digestion to remove contaminating DNA. RNA concentrations were measured by nanodrop. RNA was stored at - 80°C.

#### cDNA synthesis

cDNA synthesis was performed with the iScript Advanced cDNA synthesis Kit according to the manufacturer’s instructions with maximal shared RNA input across samples up to a maximum of 500 ng RNA template. cDNA was stored at −20°C.

#### RT-qPCR

All primer sequences are listed in Table S1. RT-qPCR reactions were performed in 10 uL total volume containing: 5 uL 2X SsoAdvanced Universal SYBR Green maser mix, 0.5 uL of 5 uM forward and reverse primer each, and 4 uL 4X diluted cDNA in nuclease-free water. Each sample was measured in technical duplicates. Amplification was measured on a LightCycler 480 system using the following cycling conditions: 1 cycle at 98°C for 5 min, 40 cycles at 95°C for 10 s, 60°C for 10 s, and 72°C for 10 s, followed by melting curve analysis. Quantitation cycles (Cq) were normalized to three housekeeping genes using geometric averaging(40). Stability of the housekeeping genes across samples was assessed using the geNorm algorithm as implemented in qbase+.

### Differential gene expression analysis

#### Library preparation and RNA-sequencing

Total RNA libraries were prepared from 8 uL purified RNA using the SMARTer Stranded Total RNA-seq Kit v3 – Pico Input Mammalian following the manufacturer’s instructions or TruSeq Stranded mRNA Library Prep Kit LT. RNA was fragmented at 94°C for 4 min prior to first strand cDNA synthesis. Library amplification was performed with 12 PCR cycles. Library size analysis was performed by running the libraries on a fragment analyzer. Library quantification was performed by RT-qPCR with the KAPA Library Quantification kit and libraries were pooled equimolarly. The final pool was quantified by Qubit. 675 pM of PicoV3-prepped libraries was loaded on a NextSeq 2000 instrument (NextSeq 2000 P2 v3, 2X 50 cycles, paired-end sequencing) with 2% PhiX or 2 pM of TruSeq-prepped libraries was loaded on a NextSeq 500 instrument (NextSeq 500/550 High-Output kit, 75 cycles, single-end sequencing) with 1% PhiX.

#### Data processing

Reads were quality controlled using FastQC (v0.11.8) and trimmed with cutadapt (v1.18) to remove any low quality bases and Illumina adapter sequences. Trimmed reads were aligned using STAR (v2.6.0) to the human hg19 reference genome assembly (GRCh37) followed by read quantification using HTSeq (v0.6.1). Count data was imported in R, and differential analysis was performed with DESeq2 (v1.42.0) with the ASO treatment as the only design variable (design = ∼ ASO). Replicate clustering was assessed by PCA, confirming clustering per condition. DESeq2-normalized counts of NESPR-targeting ASOs were contrasted to NCA to calculate a log2FC per gene. Statistical significance was determined using the default DESeq2-implemented Wald test, and a ≤ 5% Benjamini-Hochberg-adjusted P-value cutoff used to support significance. Genes with a baseMean of ≤ 10 were removed. A log2FC cutoff of ζ −0.25 and σ; 0.25 was used to filter background contrasts. Experiments were performed per cell line, and log2FCs per cell line were combined per gene. Consistent NESPR regulated genes were determined by filtering based on statistical significance and the directionality of expression changes across cell lines. Heatmaps were generated using ComplexHeatmap (v2.18.0) in R. ADRN and MES gene signatures are from van Groningen et al. (7) and were supplemented with mTFs identified by Thirant et al. (18), IEG gene signature is from Arner et al. (41), PN gene signature is derived from the Human PN Annotation effort (second version, release data 2024_0414, data accessed on 5^th^ of December 2024) of the Proteostasis Consortium (https://www.proteostasisconsortium.com/pn-annotation/), ISR gene signature is from Wong et al. (42), DDR gene signature from Knijnenburg et al. (43), and OSR based on Gene Ontology (GO:0006979). Genes present in at least three different cancers of the TSGene 2.0 database(44) were considered tumor suppressive genes. To this list, HEXIM1(39,45–47) and MIR22HG(48–51) were added as tumor suppressive genes based on available literature.

#### iRegulon analysis

Transcription factor motif enrichment scoring was performed with iRegulon (v1.3) within Cytoscape (v3.10.2). Up- and downregulated genes in the NESPR gene signature were used as separate input files. A region of 10 kb up- and downstream (total of 20 kb) of the TSS was set as the search space. The lists were queried against the 10K motif collection (9713 PWMs) and ENCODE ChIP-seq tracks (1120 experiments). A minimum NES score of 3.0, and maximum FDR for motif similarity of 0.001 were used.

#### Gene set enrichment analysis

Differential expression data sets were pre-ranked by log2FC including all baseMean-filtered genes. GSEA analyses were ran with 10 000 permutations and an FDR-based multiple testing adjustment to assess statistical significance. A ≤ 5% cutoff was considered to support statistical significance. GMT files for IEG, ADRN, and MES gene signatures were custom-made based on Arner et al. (41) and van Groningen et al. (7) supplemented with mTFs from Thirant et al. (18), respectively. Analyses and visualization were performed with the clusterProfiler (v4.10.0) R package(52). NES values were calculated as the maximum absolute running enrichment score for each separate analysis, with directionality based on the directionality of the log2FC for that specific gene.

### Genome topology

#### Preparation of 4C templates and sequencing libraries

The 4C templates were prepared as described previously van de Werken et al. (53). In brief, 10 x 10^6^ SH-SY5Y or SK-N-BE(2)c cells were seeded per template. The day after seeding or 48 hours after ASO transfection cells were detached by trypsinization, and cross-linked with 2% formaldehyde for 10 min at room temperature. Cell lysates were digested with 400 U DpnII, and proximal DNA fragments were ligated with 50 U T4 DNA ligase. Ligated DNA circles were de-crosslinked overnight using proteinase K and purified with NucleoMag P-Beads to obtain an intermediate 3C template. 3C templates were digested with 50 U Csp6I and religated as described before, resulting in 4C templates. Adaptor-containing reading and non-reading primers, specific to the viewpoints of interest, were designed to amplify interacting genomic regions relative to the viewpoint. For each viewpoint, 16 PCR reactions, with 200 ng 4C template as input, were pooled. Resulting 4C-sequencing libraries were purified using QIAquick PCR Purification kit.

#### 4C-sequencing and data analysis

4C-sequencing libraries were pooled, and 1.6 pM with 1% PhiX was loaded on an NextSeq 500 instrument (NextSeq 500/550 High-Output kit, 75 cycles) for single-end sequencing. Sequencing libraries were demultiplexed based on the presence of the viewpoint-specific reading primer sequence preceding the captured DNA sequence. After removal of the reading primer prefixes, captured sequences were mapped to the hg19 human reference genome assembly (GRCh37) with Bowtie2 (v2.2.6). Mapped reads on the *cis*-chromosome were binned per DpnII restriction fragment. Finally, 4C coverage profiles were obtained by per fragment normalization to reads per million (RPM) on the *cis*-chromosome and smoothing the normalized coverage use the rollmean function of the zoo (v1.8.12) R package with a window size of 21 fragments.

### Chromatin accessibility

25 0000 SK-N-BE(2)c cells were seeded, and next day transfected with ASO2, ASO10, or NCA. Experiments were performed in triplicates. Forty-eight hours post transfection cells washed once with PBS, pelleted, and genomic DNA was extracted from 90% of the pellet. The remaining 10% was used to confirm NESPR knockdown after ASO treatment prior to proceeding with downstream sample processing. Genomic DNA was tagmented by Tn5 transposase and DNA libraries were generated as described previously(54). Libraries were quantified by RT-qPCR with the KAPA Library Quantification kit and were pooled equimolarly.

1.9 pM of the final pool and 1% PhiX were loaded, and paired-end sequencing was performed on a NextSeq 500 instrument (NextSeq 500/550 High-Output kit, 150 cycles) with a 150 bp read length. Sequencing quality was verified with FastQC, reads were trimmed with NGMerge, alignment to the hg38 human genome assembly (GRCh38) was performed with Bowtie2 (v2.2.6), and duplicate reads were removed with SAMTools. MACS2 (v2.1.2) was used for peak calling. Peaks with an adjusted P-value ≤ 5% were considered significant, and significant peaks with a 50% overlap across samples were merged into a single matrix. Reads were counted with the SummarizeOverlaps function from the GenomicAlignments (v1.38.0) R package with the overlap matrix and corresponding BAM files per replicate and condition as input. Read counts per called overlapping peak were used as input for DESeq2 differential analysis, similar as described above (design = ∼ ASO). Contrasts of NESPR-targeting ASO compared to NCA were overlapped to identify differential open chromatin regions in both ASO treatments. These regions were filtered based on log2FC directionality and a ≤ 5% adjusted P-value significance threshold. BED files of the shared significant up- and downregulated regions were generated in R, on which the findMotifsGenome function of Homer (v4.10.3) was performed for motif enrichment analysis.

### Shotgun proteomics

#### Sample preparation and LC-MS/MS analysis

1.2 x 10^6^ IMR32 cells were seeded in quadruplicates per ASO treatment, and 48 hours later transfected with ASO1, ASO2, or NCA. Cells were washed once with PBS, collected by scraping, snap-frozen dry, and stored at −80°C. RNA was extracted from 30% of the samples for RNA as described above for RT-qPCR to confirm NESPR knockdown and subsequent RNA-sequencing. The remaining 70% was resuspended in SDS lysis buffer, containing 5% SDS, 50 mM TEAB (pH 8.0) to promote lysis. Proteins in the lysate were reduced and alkylated with 10 mM TCEP and 40 mM chloroacetamide, respectively. Samples were incubated at 95°C for 10 min with agitation. Proteins were acidified to a final concentration of 2.5% (v/v) phosphoric acid (PA), and mixed with 100 mM TEAB in 90% MeOH, and loaded onto S-Trap^TM^ Micro columns. Column-retained proteins were washed four times on the S-Trap with 150 uL 100 mM TEAB in 90% MeOH. An overnight on-column trypsin was performed using 2 ug trypsin in 20 uL 50 mM TEAB at pH 8.0 at 37°C. Next day, peptides were sequentially eluted with 50 mM TEAB, 0.2% formic acid (FA), and 50% acetonitrile (ACN). Peptides were purified on Omix C18 tips and dried in a speed-vac.

Purified peptides were redissolved in 30 uL solvent A (98% CAN, 0.1% TFA) of which 15 uL was injected on an Ultimate 3000 RSLCnano system in line connected to a Q-Exactive Biopharma mass spectrometer equipped with a Nanospray Flex Ion source. Trapping was performed at 20 uL/min for 2 min in loading solvent A on a 5 mm trapping column. Peptide separation was performed on a 250 mm Aurora Ultimate at a constant temperature of 45°C. Peptides were eluted with a non-linear gradient going from 1% solvent B (80% ACN, 0.1% TFA) reaching 33% solvent B in 60 min, 55% in 75 min, 70% in 90 min, followed by a washing step with 70% solvent B for 10 min and re-equilibration with solvent A. The mass spectrometer was operated in a data-independent mode, automatically switching between MS1 and MS2 acquisition. Full-scan spectra (375 - 1500 *m/z*) with a target value of 5 x 10^6^ at a resolution of 50 000. Precursor isolation was performed by 30 quadrupole isolations at a width of 10 *m/z* for HCD fragmentation at a normalized collision energy of 30%. MS2 spectra were acquired at a resolution of 15 000 at 200 *m/z* without multiplexing. Samples were run in a randomized-block: the first replicate of each condition was analyzed first, followed by the second replicate of each condition, until all replicates were processed.

#### Database search and differential protein expression analysis

All RAW files were searched together with the DIA-NN algorithm (v1.8.1) with default search settings, including an FDR set at 1% on precursor and protein levels. Spectra were searched against the human proteome (SwissProt database, version of August 2023), containing 20423 proteins. Enzyme specificity was set as C-terminal to arginine and lysine, also allowing cleavage at proline bonds with a maximum of two missed cleavages. Variable modifications were set to oxidation of methionine residues and acetylation of protein N-termini with a maximum of 5 variable modifications. Cysteine carbamidomethylation was set as a fixed modification. Most default settings were used, with the exception of a precursor mass range filter of 400 – 900 *m/z* and a minimal precursor charge of +2.

Proteins were quantified using DIA-NN’s implementation of the MaxLFQ algorithm, calculating protein intensities based on peptide-level quantification while accounting for shared peptides and normalization of label-free quantification (LFQ) intensities across samples. PG.MaxLFQ values were log2-transformed, and the run order was implemented as a batch effect using the removeBatchEffect function in the limma (v3.58.1) R package. Replicate clustering was assessed by PCA, confirming clustering per condition. Proteins only identified *N*-1 replicates in either the NESPR-targeting ASO or NCA were retained for downstream statistical analysis, missing values were imputed by quantile regression with imputeLCMD (v2.1) R package. Statistical analysis was performed with limma. A linear model was fitted onto the data and differential analysis was performed by empirical Bayes moderated t-tests with a Benjamini-Hochberg correction for multiple testing. Proteins with an adjusted *P-*value ≤ 5% were considered significant.

### Zebrafish

Zebrafish were housed in a Zebtac semiclosed recirculation housing system (Techniplast, Italy) and maintained following standard procedures. All zebrafish studies and maintenance of the animals were in accordance with protocol #17/100, approved by the Ethics Committee for the use of animals in research of the Faculty of Medicine and Health Sciences, Ghent University. Zebrafish were all of the AB background strain. The Tg(dβh:EGFP; dβh:MYCN) line (TgMYCN_TT)(55) and the *nespr* TSS-deletion line(56) were kind gifts from Thomas Look (DFCI, Boston, USA) and Alexander Schier (Biozentrum Universität Basel, Basel, Switzerland), respectively. Lines were genotyped as described before in the respective publications(55,56).

Heterozygous *nespr* TSS-deleted fish (designated as nespr^+/-^) were first crossed with Tg(dβh:EGFP; dβh:MYCN) to generate stable [Tg(dβh:EGFP; dβh:MYCN); *nespr*^+/-^] fish. Subsequently, these fish were crossed with *nespr*^+/-^ fish. At 5 days post fertilization (dpf), the fish were screened for EGFP expression, reflecting MYCN-expressing fish, using a fluorescence microscope with the NIS Elements software package. Fish lacking EGFP expression were excluded from downstream experiments. At 12 weeks post-fertilization (wpf), fish that were EGFP+ at 5 dpf were genotyped for *nespr* as described previously(56), and divided into three groups based on the genotype (*nespr*^+/+^, *nespr*^+/-^, *nespr*^-/-^). These groups were screened for EGFP+ tumors with a fluorescence microscope, and tumor penetrance per genotype was determined based on the final contingency table.

### Survival and correlation analyses

#### Ranksum analysis and lncRNA expression correlation with the ADRN gene signature

For each NB cell line in the RNA Atlas, the cumulative expression of the ADRN signature was computed by summing the normalized expression levels (log2[TPM + 1]) of all ADRN signature genes. NB cell lines were ranked in descending order (more ADRN to less ADRN) based on this cumulative expression. Pearson’s correlation was calculated between the ranked cumulative signature values and the expression levels of each expressed lncRNA across all cell lines. *P*-values were adjusted with Benjamini-Hochberg to control for multiple testing. Correlation coefficients (*r*) and associated log10(-*P*-values) were plotted to visualize significant (*P* ≤ 5%) associations.

#### Survival analysis

Survival analysis was performed with both overall survival (OS) and event-free survival (EFS) across different NESPR classification groups in the SEQC NB patient cohort. Classification was based on the median NESPR expression level across all patients, with expression levels higher or lower than the median expression designated as ‘high’ or ‘low’, respectively. OS represents months from diagnosis to the occurrence of patient death or censoring. EFS represents months from diagnosis to either tumor progression, recurrence, death, or censoring. Survival probabilities were calculated using the survfit function of the survival (v3.7.0) R package. Plots were generated with the ggsurveplot function of the survminer (v0.4.9) R package. The log-rank test was used to compare survival distributions between NESPR classification groups, *P*-values ≤ 5% were considered significant.

Cox proportional hazards regression models were used to perform a multivariate survival analysis to assess NESPR’s prognostic value independent from established risk markers INSS staging, age of diagnosis, and MYCN amplification status. Predictor selection was performed using a stepwise backward selection approach, starting with a full model containing all predictors and stepwise comparison with a reduced model by removing one predictor. Likelihood ratio tests (LRT) were used to assess the impact of removing each predictor on the overall model fit. If removing a predictor led to a statistically significant decrease in model fit (*P*-value ≤ 5%), the predictor was retained in the final model. Predictors of reduced models were re-evaluated with all lower-order reduced models, and this process was repeated until removing any of the remaining predictors did not lead to a significant decrease in model fit. Individual prognostic value of each predictor in the final model fit was evaluated with Wald tests (*P*-value ≤ 5%). Cox regression analyses were performed using the coxph function of the survival (v3.7.0) R package. Statistical output of the final model was extracted using the tidy function of the broom (v1.0.4) R package, with the exponentiate = TRUE and conf.int = TRUE arguments to convert log hazard ratios to Hazard ratios and the corresponding 95% confidence intervals. Forest plots were generated with ggplot2.

#### CCLE gene signature scoring

ADRN and MES signature scoring of NB cell lines in the CCLE database was conducted as described by van Groningen et al. (7). Briefly,: for each cell line, all expressed genes were ranked in descending order based on their expression (log2[TPM + 1]). The rank order of each gene was then converted to percentiles relative to the total number of expressed genes. The ADRN and MES signature scores for each cell line were calculated by averaging the percentiles of all ADRN or MES genes, respectively. Cell identities for cell lines of unknown identity were inferred by their co-clustering with cell lines of known identity based on the ADRN vs. MES signature score plot.

#### Co-expression analysis

PHOX2B and NESPR expression (log2[TPM + 1]) were correlated over all patients or cell lines in the SEQC and Westermann patient cohorts, or RNA Atlas and CCLE cell line compendia, respectively. Pearson’s correlation analysis was run with the cor.test function implemented in base R. Correlations with *P*-values ≤ 5% were considered significant, with correlation coefficients (*r*) showing the extent and directionality of the correlation.

### Conservation analysis

Nucleotide level and region level conservation of the chr4: 41,685,000 – 41,962,589 region of the hg19 human reference genome assembly (GRCh37) was assessed by visualizing the conservation scores of the UCSC Genome Browser data for PhyloP 100-way vertebrate alignment (phyloP100wayAll) and phastCons 100-way vertebrate alignment (phastCons100way), respectively. Conservation tracks were generated using the Gviz (v1.46.1) package in R, with phyloP and phastCons loaded as DataTrack objects.

### Software and statistics

All figure panels and the associated data analyses were generated and performed in RStudio with R (v4.3.2). Genomic tracks were built with Gviz (v1.46.1) with either BAM, BigWig or Bedgraph files as input for coverage plots, or BEDPE files for HiChIP genomic interaction plots using the GenomcInteractions (v1.36.0) R package compatible with Gviz. Software for specific analyses or visualizations are described above for the corresponding experiments.

For all experiments, a type I error rate (*P*-value) cutoff of ≤ 5% was considered statistically significant. Where applicable, multiple testing correction was performed using rank-based Benjamini-Hochberg *P*-value adjustment. Statistical testing of INSS staging and MYCN amplification status was performed with a Wilcoxon Rank-Sum test. Other statistical analyses used throughout the paper are specified in their designated analyses workflows described above.

## RESULTS

### NESPR is exclusively expressed in cells of the sympathetic nervous system

To identify lncRNAs associated with the ADRN CRC, we collapsed the ADRN gene signature(7) into a cumulative ADRN score based on expression data in 11 NB cell lines (8 ADRN, 1 MES, and 2 mixed), available in the RNA Atlas(29). We then correlated the expression of 2111 lncRNAs with the ADRN score across all cell lines (Fig. 1A). To substantiate the approach, we included PHOX2B, HAND2, ISL1, and GATA3 as ADRN mTFs, and PRRX1 and NOTCH2 as MES mTFs. All ADRN mTFs were significantly correlated with the ADRN score, while both MES mTFs were anticorrelated, in line with our expectations. Out of the top 10 ADRN-associated lncRNAs 5 were previously shown to be differentially upregulated in ADRN tumors compared to MES tumors(28) (Table S2), including MIAT which was previously reported as a MYCN regulator(24,57). In addition, the top ranking anticorrelated lncRNAs MIR4435-2HG and SNHG9 were previously reported to be differentially upregulated in MES tumors(28). From the top 10 ADRN-associated lncRNAs, we prioritized lncRNAs based on the tissue specificity score (tau) calculated by Modi et al. (28) in a pan-pediatric cancer study (Table S2). From the top ranking ADRN-associated lncRNAs, LINC00682 (hereafter NESPR) was the most NB-specific lncRNA. To corroborate on this, we analyzed NESPR expression in the TCGA patient cohorts comprising 10011 primary tumors over 34 different cancer types. As NB is not represented in TCGA, we integrated two different NB patient cohorts (498 and 579 tumors). Consistent with the tissue specificity scoring, we found NESPR to be uniquely expressed in tumors of the peripheral sympathetic nervous system (Fig. 1B). In addition to NB, NESPR was highly expressed in pheochromocytomas and paragangliomas (PCPG), tumors originating from the same cellular lineage as NB(58). These results were also reflected in two independent cell line collections (Fig. 1C), with the highest expression in NB cell lines. Interestingly, we observed no expression in two healthy primary human neural crest (hNCC) cell lines(9,59), suggesting NESPR expression is unique for cells committed to the SA lineage. In line with this, the NESPR expression profile in sympathoadrenal precursors (SAP)(32) demonstrates that NESPR expression is initiated once truncal NCC (tNCCs) start differentiating into SAPs, with no prior observable expression (Fig. 1D). NESPR expression seems unique for the sympathetic nervous system (SNS) developmental trajectory, as no observable expression was found in healthy mature tissues, or in terminally differentiated fetal adrenal cortices(33) (Fig. 1E). In contrast, neuroblast clusters isolated from the same fetal adrenal glands(33) demonstrated a high NESPR expression (Fig. 1E). In line with our findings, Iyer et al. (60) highlighted NESPR as a potential cell fate regulatory lncRNA during the specification of visceral motor neurons, another cell type of the SNS. Together, these data suggest that during normal development NESPR expression is confined to the SNS differentiation trajectory, and its expression is downregulated following terminal differentiation.

**Figure 1:**
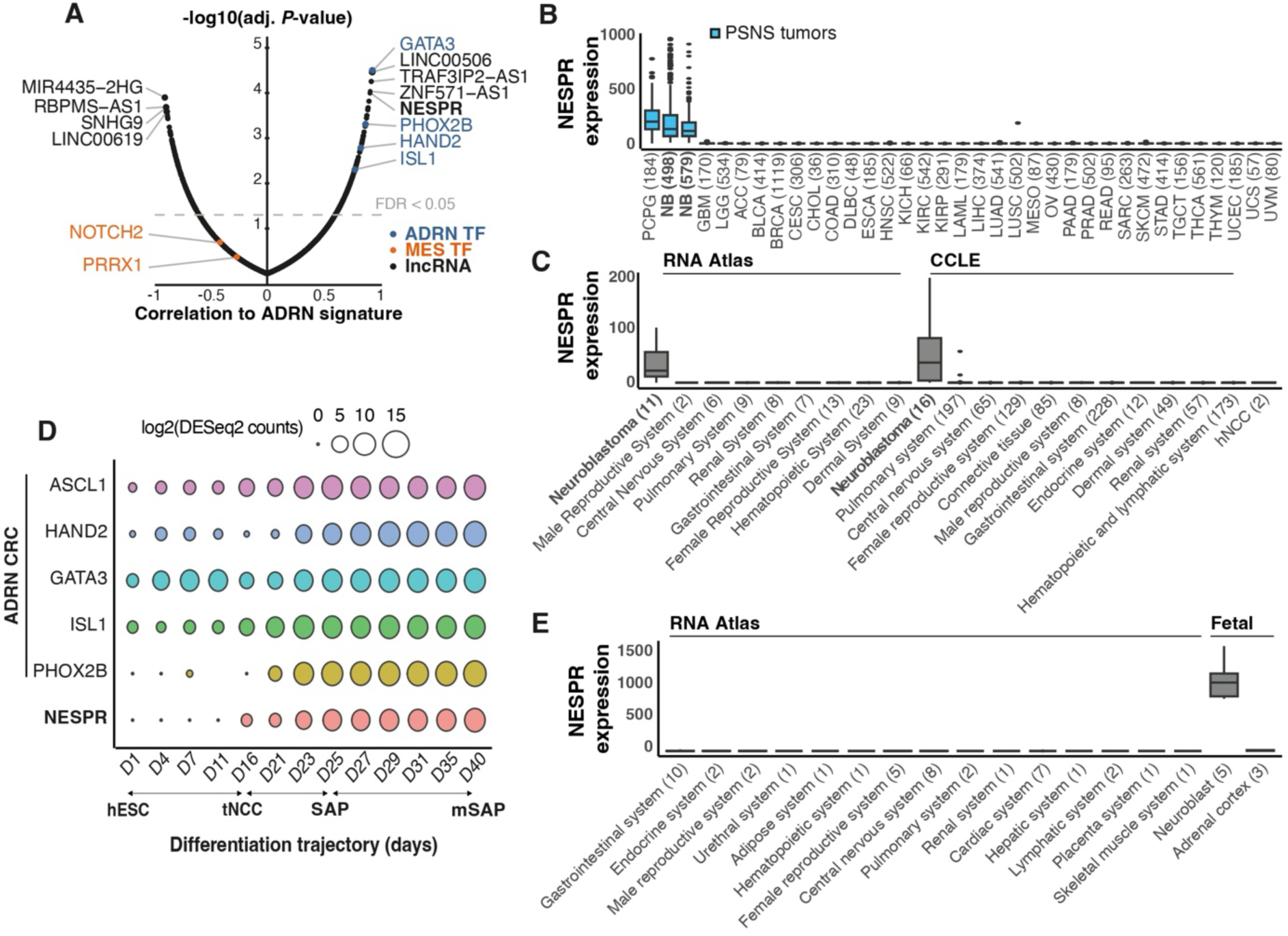
NESPR is a lncRNA specifically expressed in the sympathoadrenal lineage. (A) Correlation of expression with the ADRN gene signature of 2111 lncRNAs in a pannel of 11 NB cell lines. ADRN mTFs shown in blue, MES mTFs shown in orange, lncRNAs shown in black., top 4 correlated and anti-correlated lncRNAs are shown with NESPR in bold. FDR cutoff of 5% is shown. (B) NESPR expression profile (TPM) in 36 different patient cohorts (2 NB and 34 TCGA cohorts). Boxplots represent median expression in each cohort and are ranked by decreasing median expression. Tumors of the PSNS are shown in blue. (C) NESPR expression profile (TPM) in RNA Atlas, CCLE cell line collections, and two hNCC primary cell lines. Cell lines are grouped by human organ system. Grouped NB cell lines are shown in bold. Boxplots represent median expression in each group and are ranked by decreasing median expression. (D) NESPR expression profile (log2(DESeq2 counts)) in a SAP differentiation trajectory. ADRN mTFs are shown as a reference. (E) NESPR expression profile (TPM) in mature healthy tissues, fetal neuroblasts, and fetal adrenal cortices. Tissues are grouped per organ system. Boxplots represent median expression in each group. ADRN, adrenergic; MES, mesenchymal; TF, transcription factor; PSNS, peripheral sympathetic nervous system; hNCC, human neural crest cells;CRC, core transcriptional regulatory circuitry; hESC, humand embryonic cells; tNCC, truncal neural crest cells; SAP, sympathoadrenal precursor; mSAP, mature sympathoadrenal precursor.

### NESPR is a clinically relevant lncRNA that is expressed in preclinical NB models

As NESPR expression in tumors is restricted to the SA lineage, we assessed if NESPR expression is associated to NB patient outcomes. Kaplan-Meier analyses for overall (OS) and event-free (EFS) survival probability in the SEQC patient cohort(31) demonstrated that patients with higher than median expression of NESPR had an adverse OS and EFS compared to patients with lower expression levels (Fig. S1A). Comparison of NESPR expression between International Neuroblastoma Staging System (INSS) tumor stages and MYCN amplification status demonstrated significantly higher NESPR expression in INSS stage 4 and MYCN amplified (MNA) tumors, in two independent NB cohorts (Fig. S1B-C). A multivariate Cox regression with established risk factors (INSS, MNA, and age of diagnosis) suggests that NESPR expression serves as an independent prognostic predictor for risk assessment in NB patients (Fig. S1D). These data suggest that NESPR may have clinical relevance in NB.

While most lncRNAs are poorly conserved, NESPR shows high conservation based on both nucleotide and region level metrics (Fig. S2A). In mouse and zebrafish, two model organisms for which clinical NB models are available, we identified a syntenic annotated lncRNA gene downstream of *Phox2b/phox2bb* (Fig. S2B). Expression of the syntenic lncRNA was confirmed in LSL-MYCN and Th-MYCN NB mouse models, as well as in the *Tg(dβh:EGFP-MYCN)* NB zebrafish model (Fig. S2C).

### *NESPR* and *PHOX2B* are co-expressed from the same insulated gene neighborhood

NESPR is located to 125 kb downstream of the ADRN mTF PHOX2B. H3K27ac marks and annotated SE regions in two ADRN cell lines (IMR32 and SK-N-BE(2)c) show *NESPR* is located in the main SE regulating PHOX2B expression in these cell lines (Fig. 2A). Based on RNA-sequencing data and CAGE-sequencing profiles, we determined the main NESPR transcript is 555 nucleotides, transcribed from two exons (Fig. 2B). In line with NESPR’s ADRN-specific expression profile, H3K27ac marks and expression were absent in MES cell lines (SHEP and GIMEN; Fig. 2A-B). H3K4 methylation profiles revealed H3K4me1 marks across the *NESPR* locus, consistent with enhancer activity, while broad H3K4me3 peaks were mainly located at exonic regions of the different NESPR isoforms, a pattern that has been observed for other cell identity genes(61) (Fig. 2B).

**Figure 2:**
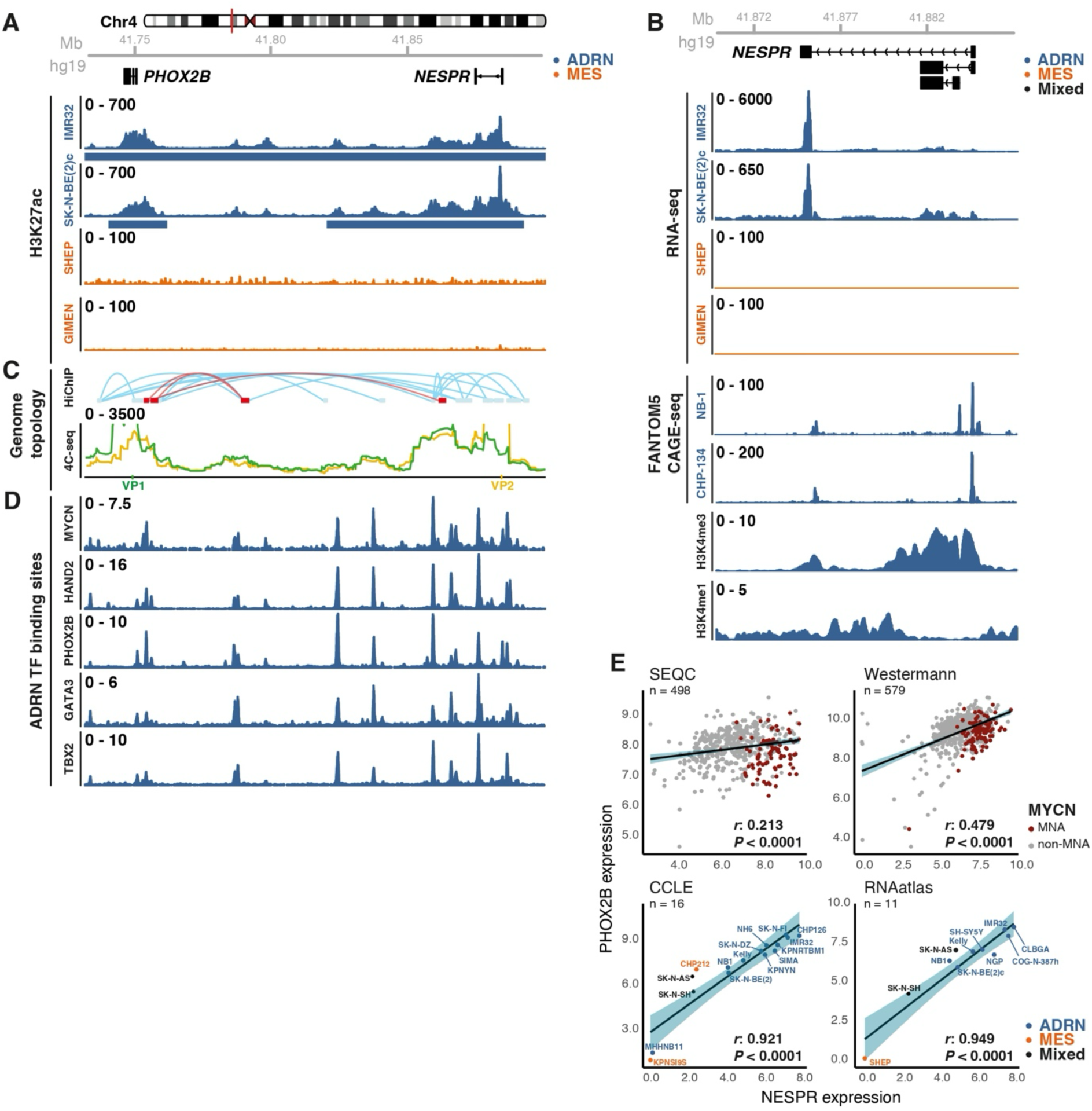
NESPR and PHOX2B form a insulated gene neighborhood. (A) H3K27ac profiles in 2 ADRN (SK-N-BE(2)c and IMR32) and 2 MES (SHEP and GIMEN) cell lines. Called SEs are shown by blue or orange bars beneath each track for ADRN and MES cells respectively. (B) Coverage tracks in NESPR locus for RNA-seq in 2 ADRN and 2 MES cell lines, CAGE-seq (FANTOM5) in 2 ADRN cell lines, and H3K4 mono- and trimethylation profiles in SK-N-BE(2)c. (C) Genome topology in the PHOX2B-NESPR genomic locus by 4C-sequencing using a PHOX2B VP (VP1) and a NESPR VP (VP2) in SH-SY5Y cells. Genomic interaction plots of SMC1 HiChIP in Kelly and CLBGA are shown on top in light blue and red, respectively. (D) ADRN mTF binding sites in SK-N-BE(2)c. (E) Co-expression of PHOX2B and NESPR in 2 NB patient cohorts (SEQC and Westermann) and 2 cell lines collections (RNA Atlas and CCLE). ADRN, adrenergic; MES, mesenchymal; TF, transcription factor.

Given the presumed gene regulatory activity of the SE region, we interrogated genomic interactions in the *PHOX2B*-*NESPR* locus using 4C-sequencing analysis in SH-SY5Y with viewpoints (VP) located at both genes. High coverage was observed at *NESPR* within the PHOX2B VP (VP1) and at *PHOX2B* within the *NESPR* VP (VP2), indicating genomic interactions occurring between both loci (Fig. 2C). Moreover, profiles between the VPs show a remarkable overlap at the same intergenic genomic regions. In support of our 4C-sequencing experiments, SMC1 HiChIP data from two independent studies in two different ADRN NB cell lines(34,35) identified the same genomic interactions between *NESPR* and *PHOX2B* as well as those observed in the intergenic region. These findings demonstrate extensive looping between the *PHOX2B* and *NESPR* genes, illustrative of a putative insulated gene neighborhood. ADRN mTF ChIP-sequencing data(8) supports co-regulation of both genes, with mTF bound regions overlapping the anchor points seen in the genome topology (Fig. 2D). In addition, these anchor points are conserved elements, suggesting genome topology and locus regulatory mechanisms are conserved as well (Fig. S2A). We evaluated co-expression of *PHOX2B* and *NESPR* in two NB patient cohorts and the RNA Atlas and CCLE cell line collections. Both NB cohorts displayed a significant co-expression, with overall higher expression of both genes in MNA cases (Fig. 2E). While the cellular identity of RNA Atlas NB cell lines is known based on literature, some cell lines in CCLE have not yet been described within the context of ADRN/MES CRCs. Therefore, we scored all cell lines as described in van Groningen et al. (7) and determined identities of cell lines of unknown identity based on co-clustering (Fig. S3A). ADRN cell lines had overall higher expression of both genes compared to MES cell lines or mixed populations. Independent of cell identity, a strong and significant co-expression was found in NB cell lines in both collections (Fig. 2E).

### NESPR regulates ADRN NB cell survival and proliferation

Goudarzi et al. (56) previously demonstrated that the nespr (named lnc-phox2bb) transcript is dispensable for normal zebrafish development. To study the role of NESPR in NB cell growth *in vivo*, we crossed *nespr* TSS-deletion fish from Goudarzi et al. (56), which lack the first exon, with Tg(MYCN_TT) fish(55), which develop NB tumors with a penetrance of ∼70% (Fig. 3A). At 12 weeks post-fertilization (wpf), tumors were observed in 83.3% and 88.9% of wildtype *nespr* and heterozygous TSS-deletion fish, respectively (Fig. 3B). In contrast, only 40% of homozygous TSS-deletion fish developed tumors (Fig. 3B), indicating that nespr is haplosufficient and contributes to full tumor penetrance. We then determined if NESPR expression is also required for the proliferation and survival of human NB cells. Real-time proliferation monitoring was performed using IncuCyte live-cell imaging for IMR32, NGP, SK-N-BE(2)c, and SHEP cell lines. SK-N-BE(2)c, NGP, IMR32 were selected to represent cell lines with low, intermediate, and high NESPR expression, respectively, while SHEP is a MES cell line that does not express NESPR (Fig. S4A). ASOs targeting NESPR were designed either at the 3’ end of the transcript targeting an exon or intron (ASO2 and ASO10, respectively) or targeting the splice junction (ASO1) to avoid triggering premature RNA PolII transcriptional termination of neighboring genes(62,63) (Fig. S4B). ASO treatment caused a significant reduction in NESPR expression of 27-57% relative to a negative control ASO (NCA) depending on the ASO and cell line (Fig. S4C). Cells treated with NESPR-targeting ASOs or NCA were monitored for confluence over 96 hours post-transfection. NESPR depletion significantly reduced confluence in NGP and SK-N-BE(2)c cells compared to the NCA (Fig. 3C). IMR32, a high NESPR-expressing cell line, also showed an initial confluence reduction, but later recovered (Fig. 3C), likely due to the lower ASO-mediated NESPR depletion (Fig. S4C). Moreover, this pattern aligns with the haplosufficiency of nespr observed in heterozygous TSS-deletion fish. In contrast, SHEP, a MES-type NB cell line that does not express NESPR, showed no reduction in confluence upon NESPR depletion (Fig. 3C). These proliferative defects were accompanied by a significant increased caspase activity at 48 hours post-transfection (Fig. 3D).

**Figure 3:**
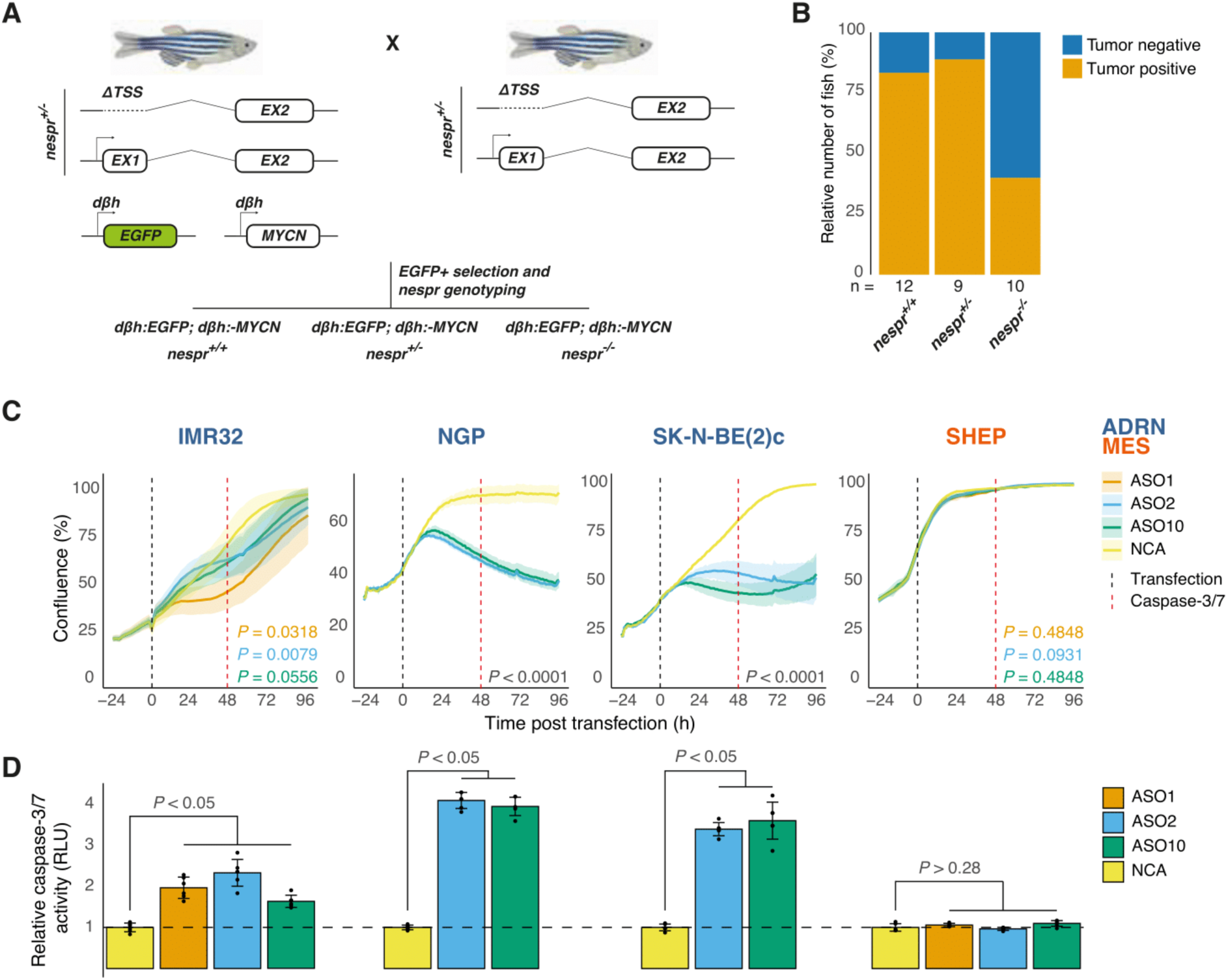
NESPR depletion impairs survival of ADRN NB cell lines. (A) Genetic crossing scheme to generate backcrossed wild type, heterozygous and homzygous nespr knock-out zebrafish in a MYCN_TT background. (B) Tumor penetrance at 12 wpf for all genotypes. (C) Proliferation of IMR32, NGP, SK-N-BE(2)c, and SHEP cell lines following ASO treatment. Black dashed line reprensents time of transfection. Red dashed line represents the time point at which caspase-3/7 experiments were performed. Time is shown relative to the time point of transfection. ADRN and MES cell lines are shown in blue and orange, respectively. (D) Relative caspase-3/7 activity in IMR32, NGP, SK-N-BE(2)c, and SHEP cells following ASO treatment. TSS, transcription start site; EX, exon; wpf, weeks post-fertilization; ASO, antisense oligonucleotide; NCA, negative control ASO; ADRN, adrenergic; MES, mesenchymal; RLU, relative light units.

### NESPR depletion triggers a stress response

To elucidate NESPR’s functionality, we depleted NESPR expression in NB cell lines and performed RNA-sequencing (Table S3). As Iyer et al. (60) previously showed PHOX2B repression after NESPR depletion, we evaluated PHOX2B levels in NB cells following NESPR depletion. While SK-N-BE(2)c and NGP showed a significant PHOX2B repression (29% and 41%, respectively), PHOX2B levels were not altered in IMR32 cells following NESPR depletion (Fig. S4D), suggesting that NESPR depletion was either not sufficient or NESPR has no regulatory impact on PHOX2B expression in IMR32 cells. Previous studies have established that enhancer-derived RNAs can modulate local enhancer-promoter looping to regulate neighboring genes *in cis*(64,65). To interrogate NESPR’s involvement in *NESPR*-*PHOX2B* looping, we performed 4C-sequencing after NESPR depletion in SK-N-BE(2)c cells but did not observe any impact on looping patterns in either VP (Fig. S4E). In addition, fractionation data in SK-N-SH(37) and SK-N-BE(2)c (Fig. 4A), and imaging of NESPR transcripts in SK-N-BE(2)c and IMR32 cells (Fig. 4B-C) demonstrated that NESPR is primarily localized to the cytosol, suggestive of a *trans* function. We overlapped all significantly up- and downregulated genes across all cell lines and ASOs into a consistent response signature with 123 up- and 175 downregulated genes (Fig. 5A). In general, ADRN signature genes were downregulated, while MES signature genes were upregulated, indicative of a molecular shift from an ADRN towards a MES identity. These observations were supported by gene set enrichment analysis using ADRN and MES gene sets (Fig. 5B). To further investigate the transcriptional response to NESPR depletion, we performed transcription factor motif enrichment using iRegulon (Table S4). Among the top-ranking TFs, SRF (Fig. 5C) stood out as a master regulator of stress response genes, specifically immediate early response genes (IEGs). Indeed, various IEGs in the NESPR response signature were upregulated (Fig. 5A), and gene set enrichment analyses further supported these observations (Fig. S4F). Moreover, SRF itself was also significantly upregulated in all differential analyses, except for ASO2-treated NGP cells. In line with these results, ATAC-sequencing after NESPR depletion in SK-N-BE(2)c cells showed motif enrichment of IEG TFs (FOS, FOSL1, JUNB, ATF3) at differential accessible chromatin after NESPR depletion (Fig. S4G). As SRF and IEGs respond to a variety of internal and external stressors, we compared the NESPR response signature to several stress response pathways: the proteostasis network (PN; includes a.o. heat shock response (HSR) and unfolded protein response (UPR) pathways), integrated stress response (ISR), DNA damage response (DDR), and oxidative stress response (OSR) pathways (Fig. 5A). We observed overrepresentation of PN genes in the NESPR response signature, indicative of impaired protein homeostasis. In line with PN deregulation, HSF1 was among the top regulators of the upregulated genes in our iRegulon analysis (Fig. 5C). Therefore, we performed differential shotgun proteomics experiments in IMR32 following NESPR depletion (Table S5). We reliably quantified 7172 proteins across all conditions, with respectively 315 and 270 significantly up- and downregulated proteins shared between ASO1 and ASO2 contrasts relative to NCA. To confirm its non-coding function, we conducted a database search that included all open reading frames (ORF) with canonical ATG start sites present in the NESPR transcript. No peptides corresponding to these ORFs were detected, further supporting the lack of any coding potential. We integrated transcriptional and proteomic data of ASO-treated IMR32 to distinguish transcriptional and translational effects (Fig. S4H). 115 and 88 genes were up- and downregulated, respectively, at both levels (Fig. 5D). Moreover, the majority of upregulated genes at both RNA and protein levels in the IMR32 NESPR response signature were PN members, suggesting a shift in proteome content or stability upon NESPR depletion. This effect was less observable in proteins that are differential only in the proteomics data (Fig. 5E). In contrast, ADRN mTFs such as GATA3, PBX3, EYA1, HAND1, SATB1, and ESRRG were downregulated only at the protein level. In addition, PHOX2B was significantly downregulated at the protein level in ASO2-treated IMR32 cells (Fig. S4I). Together these data suggest that NESPR depletion induces a stress response that causes proteomic rewiring of the ADRN CRC in favor of a more MES-like cell identity.

**Figure 4:**
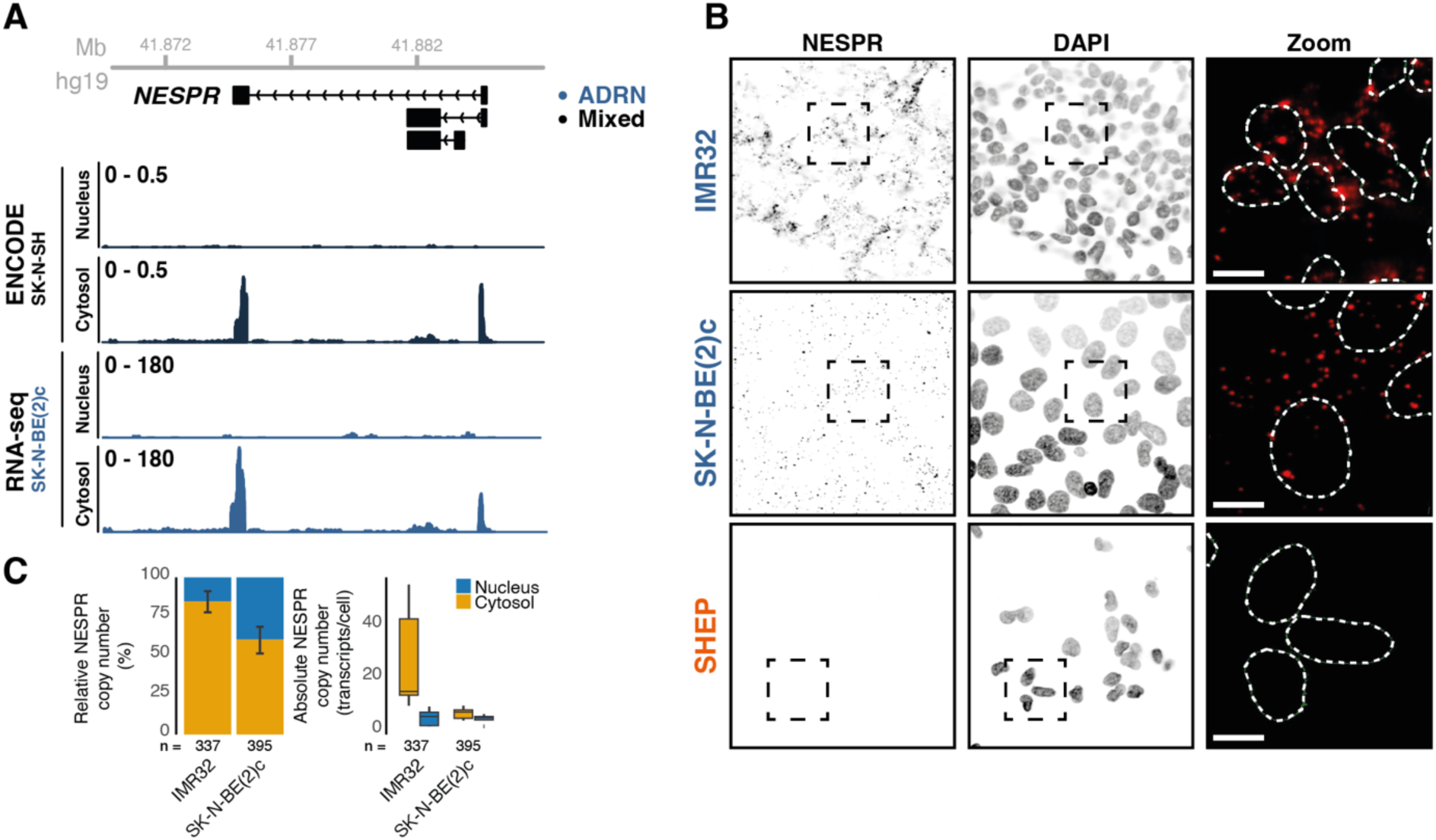
NESPR is a cytosolic lncRNA. (A) RNA-seq coverage tracks of the NESPR after cellular fractionation in SK-N-SH and SK-N-BE(2)c. (B) RNAscope of NESPR and DAPI staining in IMR32, SK-N-BE(2)c, and SHEP cells. Dashed lines represent the zoomed region. Nuclei in the zoomed region are outlined in white, NESPR is shown in red. Scales represent 5 μm. (C) Copy number quantification of NESPR in IMR32 and SK-N-BE(2)c cells. Left shows the relative number of NESPR copies in the nucleus (yellow) and cytosol (blue). Right shows the absolute number of transcripts per cell in both compartments. Boxplots represent the median number of copies. N shows the number of cells used for quantification over three different slides. ADRN, adrenergic; DAPI, 4,6-diamidina-2-phenylindole.

**Figure 5:**
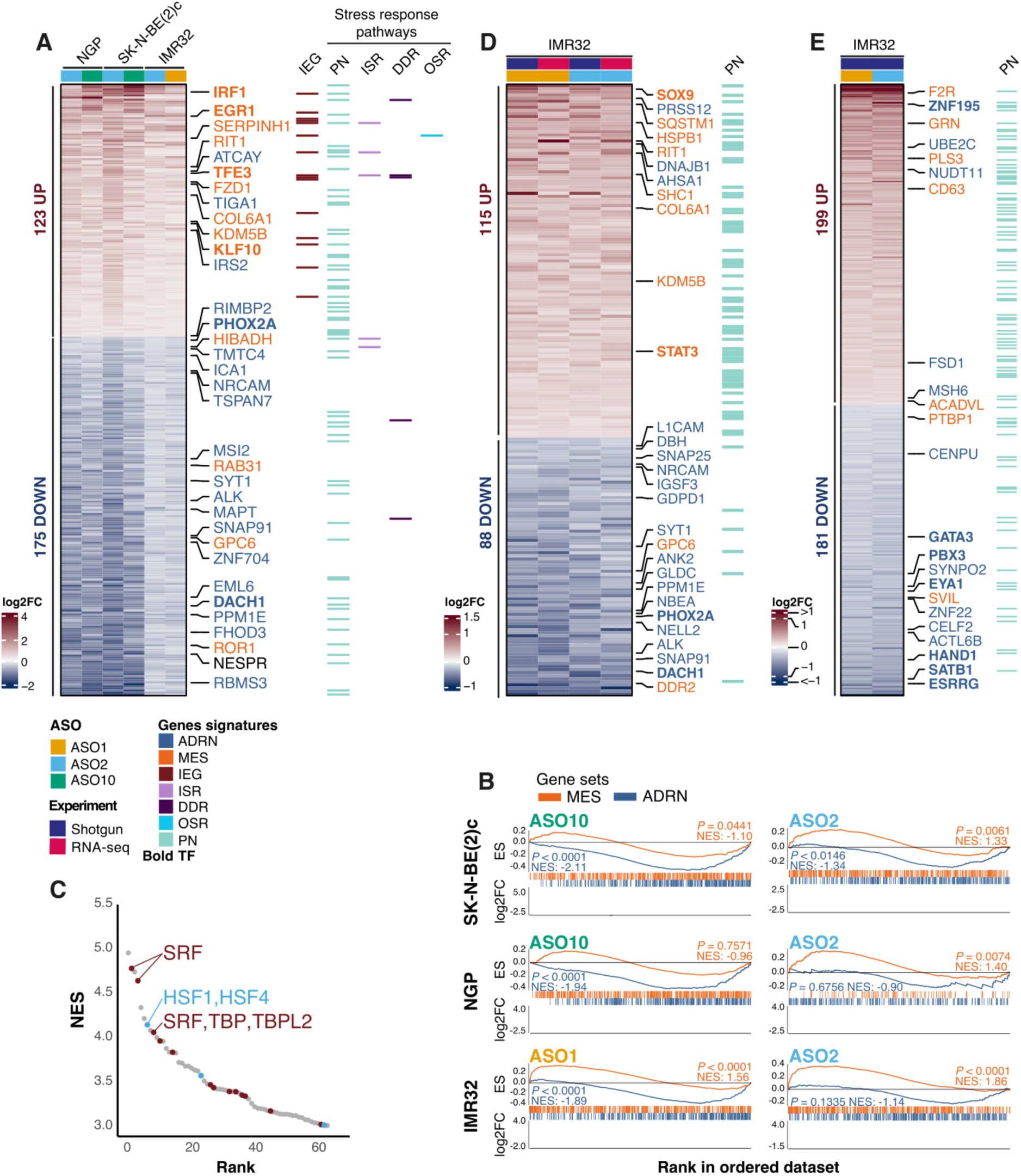
NESPR depletion induces proteostasis rewiring. (A) Transcriptional NESPR response signature in IMR32, NGP, and SK-N-BE(2)c cells. NESPR is shown in black. (B) Gene set enrichment analysis of MES and ADRN signature in all ASO experiments. (C) iRegulon output of genes upregulated in the NESPR response signature defined in (A) SRF and HSF1 annotation are shown for the top 10 ranking TFs. SRF annotated motifs are shown in brown, HSF1 annotated motifs are shown in light blue. (D) NESPR response signature in IMR32 cells that is reflected in both the RNA and protein levels. (E) NESPR response signature in IMR32 cells at the protein level only. For (A, D, E), MES and ADRN signature genes are shown and are highlighted in orange and blue, respectively. TFs are shown in bold. ADRN, adrenergic; MES, mesenchymal; ASO, antisense oligonucleotide; NCA, negative control ASO; IEG, immediate early genes; ISR, integrated stress response; DDR, DNA damage response; OSR, oxidative stress response; TF, transcription factor; PN, proteostasis network; ES, running enrichment score; NES, normalized enrichment score.

## DISCUSSION

Our present work elaborates on the involvement of the lncRNA NESPR in intratumoral heterogeneity of NB cells. We demonstrate that NESPR is strongly associated with the ADRN CRC and uniquely expressed in cells of the SA lineage, both in a developmental and disease context. Owing to its genomic context, NESPR expression is regulated by the ADRN CRC and is closely associated with PHOX2B expression, its protein-coding neighbor. We demonstrate that extensive genomic interactions occur between both genes, exemplifying an insulated gene neighborhood in which genes are co-regulated and share transcriptional resources to facilitate their expression. Our data does not support a regulatory role for the NESPR transcript in the maintenance of these genomic interactions, consistent with its primarily cytosolic location, nor does it support a role in transcriptional regulation of PHOX2B *in cis*. In contrast, we demonstrate that depletion of NESPR leads to transcriptional reprogramming, characterized by the induction of IEGs, which are commonly associated with stress responses. This stress response encompasses proteome rewiring indicated by upregulation of the proteostasis network, and ultimately leads challenged cells through a selective bottleneck associated with a shift from an ADRN to a MES-like molecular phenotype.

High-throughput sequencing has revealed extensive transcription from enhancer elements, giving rise to a diverse class of RNA molecules collectively referred to as enhancer RNAs (eRNA). Traditionally, eRNAs are characterized by bidirectionally transcription, short length, lack of processing (absence of splicing, 5’ capping, and 3’ polyadenylation), and inherent instability, making them detectable only by nascent RNA-sequencing. A considerable proportion of lncRNAs are transcribed from enhancer elements, with studies suggesting that eRNAs contribute to the emergence of lncRNA genes(66–68). This notion is further supported by studies demonstrating that certain lncRNAs are essential for enhancer activity(68). NESPR aligns with this paradigm due to its high degree of conservation, its genomic location, and its role in regulating PHOX2B in ADRN cell lines. However, Goudarzi et al. (56) demonstrated no changes in phox2bb levels in heterozygous *nespr* TSS-deletion fish, indicating that nespr does not affect enhancer activity during normal zebrafish development. This discrepancy between our *in vitro* data and those of Goudarzi et al. (56) may be attributed to differences in models systems or species, with human NESPR orthologues having an additional function absent in zebrafish. Notably, PHOX2B deregulation was observed in NGP and SK-N-BE(2)c cells but not in IMR32, suggesting that this effect may result from the collapse of the ADRN CRC rather than a direct consequence of NESPR depletion. Goudarzi et al. (56) previously demonstrated that the nespr transcript is dispensable for zebrafish development. However, when we introduced the same *nespr* TSS-deletion line into a Tg(MYCN_TT) background, we observed a decrease in tumor penetrance in *nespr*^-/-^ mutants. These findings raise the question of whether NESPR plays a role in human development and whether its function (along with that nespr) in neuroblastoma reflects a normal developmental function or represents a gain-of-function event specific to tumorigenesis. In addition, our *in vivo* experiments demonstrate that *nespr* is haplosufficient, consistent with our proliferation data in the high NESPR-expressing cell line IMR32 after NESPR depletion. Moreover, its haplosufficiency is in line with a *trans* function in the cytosol.

IEGs are rapidly and transiently induced following extracellular stimuli or stressors to initiate signaling cascades that result in either stress relief or induction of cell death. In this context, IEGs also have a crucial role in triggering cell state transitions during developmental programs(69). IEG expression has been extensively linked with receptor tyrosine kinase (RTK) activation(70), a process driven by extracellular signals that leads to downstream activation of the RAS/MAPK signaling cascade(71). Thirant et al. (18) found that EGF- and TNFα-induced RTK signalling in ADRN cells contributed to a spontaneous shift towards a MES identity. Additionally, Gartlgruber et al. (34) demonstrated RAS activity is elevated in MES tumors and is sufficient to induce cell state switching to a more MES-like phenotype *in vitro*. A recent preprint highlights EGR1 and FOS as key regulators of the RAF-MAPK signalling response(72), a pathway downstream of RAS activation. The most notable RTK described in NB is ALK. Interestingly, ALK transcript and protein levels are downregulated following NESPR depletion, suggesting that ALK signaling may not contribute to MES-associated RAS activity. This also implicates NESPR in regulating ALK signaling or in preventing RAS activity in ADRN cells. Interestingly, several IEGs are integral to the MES gene signature, including MES mTFs such as EGR1, EGR3, FOSL1, FOSL2, MAFF, ID1, and IRF1. Notably, some of these genes are upregulated following NESPR depletion, suggesting that the MES cell state is intrinsically linked to general stress responsiveness. This notion aligns with *in vitro* and xenograft studies demonstrating NB cell state switching to confer resistance to chemotherapy(9,73) or targeted treatment(16,74,75). While the overexpression of PRRX1(7) or the intracellular domain of NOTCH3(14) have been shown to induce an ADRN to MES shift, *ARID1A* knock-out(17) represent the only tested genetic deficiency demonstrated to drive this process. Here, we demonstrate that NESPR depletion causes cellular stress, leading to a molecular rewiring toward a MES molecular phenotype. Whether this cellular stress arises directly from NESPR depletion or as a consequence of an ADRN to MES shift downstream of NESPR depletion remains to be determined. Notably, while some ADRN mTFs (DACH1, PHOX2A) are transcriptionally downregulated, others (GATA3, PBX3, EYA1, HAND1, SATB, and ESRRG) are specifically downregulated (post)translationally. This suggests that NESPR mechanistically impacts either translational regulation or protein degradation dynamics, consistent with its cytosolic localization.

Cell state transitions are generally depicted as binary events. However, research on epithelial-to-mesenchymal transition (EMT) has demonstrated that these transitions occur through a spectrum of intermediate states, forming a continuum of hybrid states characterized by simultaneous expression of molecular markers from both endpoints states(76). Similarly, hybrid or transitional states have been proposed for NB(18,77). These studies(18,77) and others(28) identify NESPR as a signature gene for ADRN cells, while Thirant et al. (18) even observed NESPR expression in ADRN cells with mesenchymal features. Together, our data obtained on *in vitro* cellular models and tumor patient data identify NESPR as an ADRN-specific lncRNA that is essential for ADRN NB cell survival. We demonstrate that NESPR depletion induces molecular cell state switching. Given recent (pre)clinical advances(78,79), a broader understanding of ncRNA dependencies will widen our search space for novel druggable targets to improve patient outcomes.

## DATA AVAILABILITY

The 4C-sequencing, RNA-sequencing, and ATAC-sequencing data discussed in this study have been deposited in NCBI’s Gene Expression Omnibus and are accessible through GEO series accession numbers GSE293714, GSE293715, and GSE293716, respectively.

The mass spectrometry proteomics data have been deposited to the ProteomeXchange Consortium via the PRIDE partner repository with the dataset identifier PXD062067 (reviewer token: utaq8obh2VFt; or username: with password: x5bCWtK4UJLI).

All data will be made public upon publication.

## AUTHOR CONTRIBUTIONS

Conceptualization: L.Del., D.R., E.d.B., T.F.B, L.Dep., B.D., T.S., F.S., S.E., P.M.; Formal analysis: L.Del., D.R., E.d.B. E.D., T.F.B., L.Dep., S.V.H., G.M., R.M., L.L.D., J.A.; Funding acquisition: F.S., S.E., P.M.; Investigation: L.Del.; D.R., E.d.B., E.D., T.F.B., L.Dep., B.D., S.V.H., G.M., R.M., L.L.D., J.R., K.V., N.Y.; Methodology: L.Del.; D.R., E.d.B., E.D., T.F.B., L.Dep., S.V.H., G.M., J.R., K.V., N.Y., B.M., S.V., P.V.V., S.S.R., G.P., F.S., S.E., P.M.; Project administration: L.Del.; D.R., E.d.B., F.S., S.E., P.M.; Software: L.Del.; E.D., B.D., J.R., J.A.; Resources: S.V.H., G.M., J.R., B.M., S.V., P.V.V., S.S.R., G.P., T.S., F.S., S.E.; Supervision: T.S., F.S., S.E., P.M.; Visualization: L.Del.; Writing – original draft: L.Del., P.M.; Writing – review & editing: all authors.

## ACKNOWLEDGEMENTS

The authors acknowledge the VIB Proteomics Core, VIB Bioimaging Core Ghent, and ERA Platform Research Core (VUB) for their assistance on the presented work. Certain Figures contain elements from Biorender.

## Supplementary Data

**Fig. S1:**
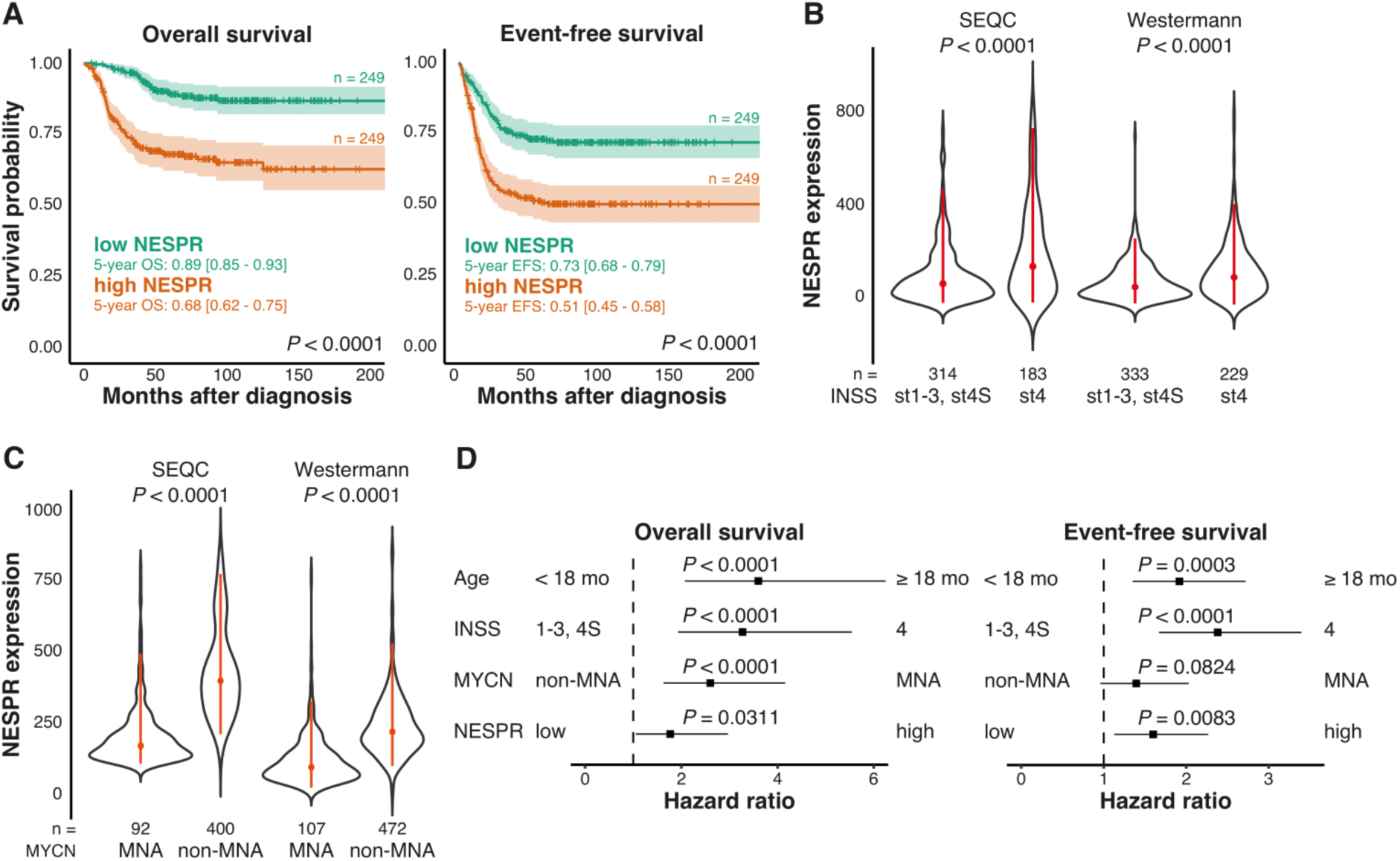
NESPR correlates with clinical parameters in high-risk neuroblastoma. (A) Kaplan-Meier analysis and 95% confidence intervals for OS and EFS in the SEQC patient cohort with grouping based on median NESPR expression across the cohort. 5-year survival rates are highlighted. (B) NESPR expression (TPM) in the SEQC and Westermann patient cohorts with grouping based on INSS tumor classification. Red lines represent 25% quantile, median, and 75% quantile values. (C) NESPR expression (TPM) in the SEQC and Westermann patient cohorts with grouping based on MYCN amplification status. Red lines represent 25% quantile, median, and 75% quantile values. (D) Multivariate Cox regression analyses for OS and EFS for known prognostic predictors and NESPR expression (median cutoff). For all panels, n represents the number of patients. OS, overall survival; EFS, event-free survival; MNA, MYCN amplified.

**Fig. S2:**
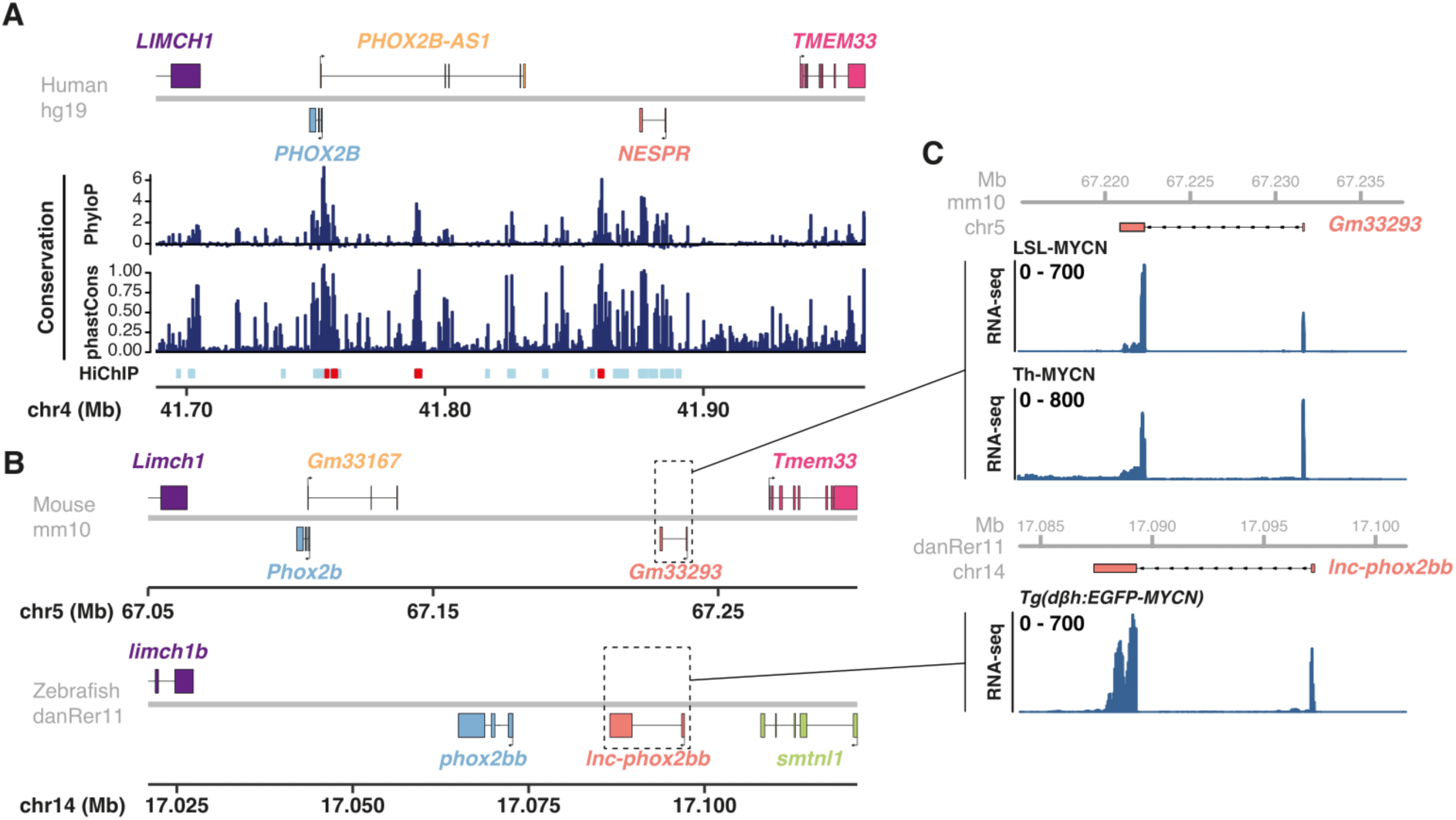
NESPR is a conserved lncRNA with expression in neuroblastoma models. (A) Conservation of the *PHOX2B-NESPR* insulated gene neighborhood in human. PhyloP and phastCons show nucleotide and region-based conservation over 100 vertebrate species. SMC1 HiChIP annotated regions in Gartguber *et al.* and Debruyne *et al*. are shown in red and light blue. (B) Syntenic regions in mouse and zebrafish. Conserved genes have the same color. (C) RNA-sequencing coverage of syntenic NESPR in mouse and zebrafish neuroblastoma models. Mb, megabases; chr, chromosome.

**Fig. S3:**
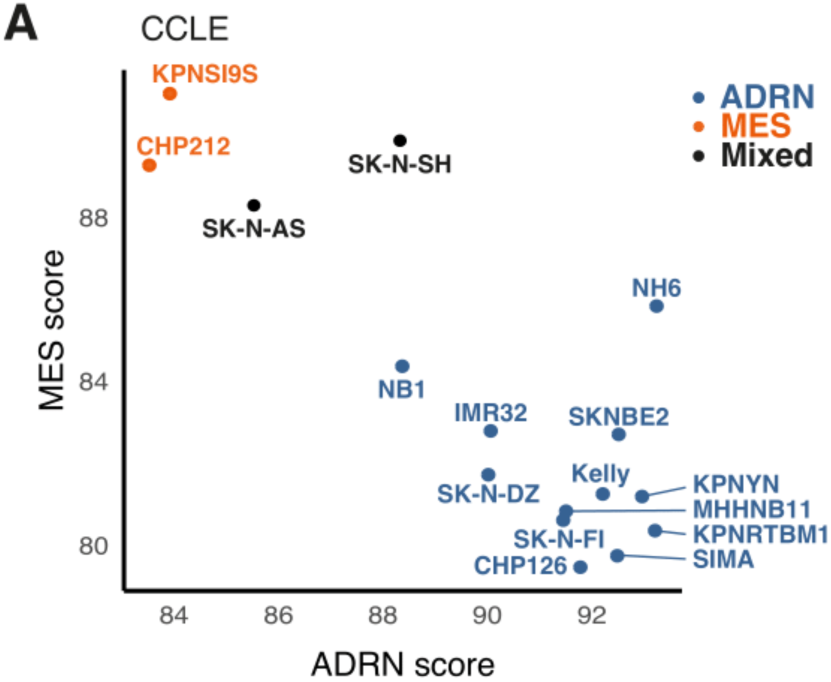
Cell identity clustering of NB cell lines in CCLE. (A) Score clustering of ADRN and MES scores in all NB cell lines in CCLE. ADRN, adrenergic; MES, mesenchymal.

**Fig. S4:**
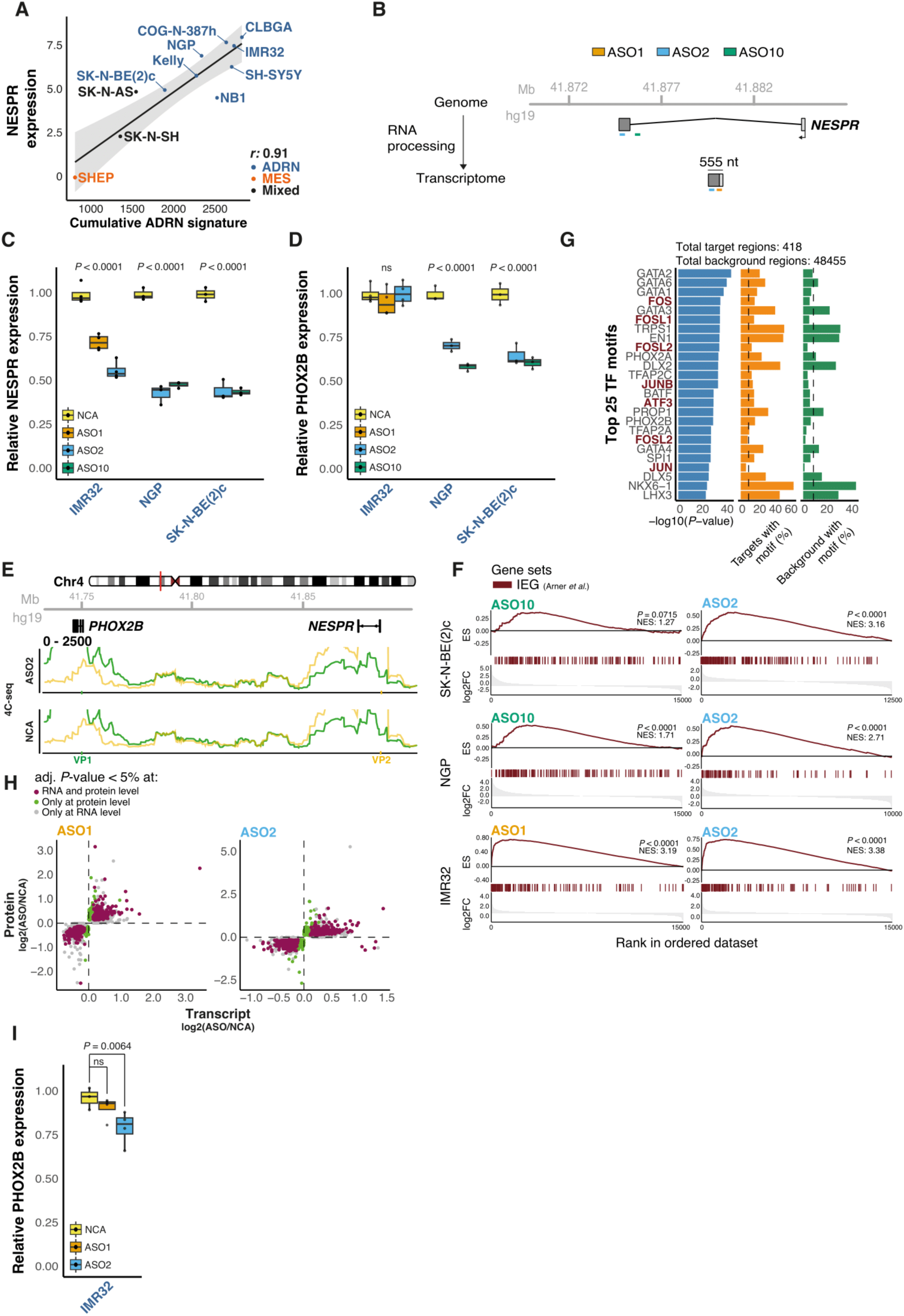
NESPR depletion induces a stress response. (A) Schematic overview of the ASO design strategy. (B) Linear regression of NESPR expression (log2(TPM+1)) and the cumulative ADRN score of all NB cell lines in RNA Atlas. Blue represent ADRN lines, black mixed populations, and orange MES lines. *r* represents the Pearson’s correlation coefficient. Light grey area represents the 95% confidence interval. (C) Relative NESPR and PHOX2B levels of ASO-treated IMR32, NGP, and SK-N-BE(2)c cells compared to a NCA. Boxplots represent median values with 25% and 75% quantiles. (D) 4C-sequencing coverage profile of ASO- and NCA-treated SK-N-BE(2)c cells at the *PHOX2B-NESPR* insulated gene neighborhood. Colors represent the PHOX2B (VP1, green) and NESPR (VP2, yellow) viewpoints. (E) Gene set enrichment analysis of IEGs in all differential RNA-sequencing experiments. (F) Motif enrichment output for the top 25 TFs in differential accessible regions after NESPR depletion. (G) Relative transcript (RNA-sequencing) and protein (shotgun proteomics) levels for each ASO treatment in IMR32 cells over NCA treatment. Genes significant at both levels and with the same directionality are shown in burgundy. Genes only significant at either the transcript and proteins level are shown in gray or green, respectively. Genes not significant in both experiments are not shown. (H) Relative PHOX2B protein levels in IMR32 cells after NESPR depletion compared to a NCA. Boxplots represent median values with 25% and 75% quantiles. ASO, antisense oligonucleotide; ADRN, adrenergic; MES, mesenchymal; TF, transcription factor; NCA, negative control ASO; IEG, immediate early gene; ES, running enrichment score; NES, normalized enrichment score; Mb, megabases; chr, chromosome.

